# Efficient treatment of a preclinical inflammatory bowel disease model with engineered bacteria

**DOI:** 10.1101/619536

**Authors:** Szilamer Ferenczi, István Horváth, Natália Szeőcs, Zsuzsanna Grózer, Dániel Kuti, Balázs Juhász, Zsuzsanna Winkler, Tibor Pankotai, Farkas Sükösd, Krisztina J. Kovács, Zoltan Szallasi

## Abstract

We developed an engineered bacterium based, RNA interference mediated therapeutic method to significantly reduce the symptoms in the most frequently used animal model of inflammatory bowel disease. This transkingdom RNA interference strategy was based on the non-pathogenic *E. coli* MDS42 strain, which was engineered to constitutively produce invasin and histeriolysin cytolysin. These proteins enabled the bacteria first to invade the colon epithelium and then degrade in the phagosome. This allowed the delivery of a plasmid encoding shRNA targeting TNF alpha into the cytoplasm of the target cells. The expression levels of TNF alpha and other cytokines significantly decreased and the inflammatory symptoms were significantly reduced. With further safety modifications this method could serve as a safe and side-effect free alternative to biologicals targeting TNF-alpha.

## Introduction

Inflammatory bowel disease (IBD) is an often severe, chronic disease of the colon and small intestine. It is characterized by recurrent inflammation of the gastrointestinal tract and have serious, potentially lethal consequences. Its exact pathomechanism is not known but increased expression of TNF-alpha plays an important role in the disease. This mechanistic insight led to the development of one of the more effective treatments of the disease, biologicals, such as infliximab, that inhibit the function of TNF-alpha^1^. This treatment option has several drawbacks that warrant the development of alternative treatments, such as high cost, severe potentially lethal side effects and the fact that up to 50% of patients do not respond or lose sensitivity to this treatment over time1.

An appealing idea is to use genetically engineered bacteria that could deliver inhibitory molecules, such as shRNA, directly into the target cells specifically against inflammatory mediators such as TNF alpha. In order to achieve this we used a non-pathogenic *Escherichia coli* MDS42 strain with significantly reduced evolvability^2^. This strain, *E.coli* MDS42, was modified to express *invasin* that allows bacteria invade mammalian cells and *listeriolysin O* cytolisyn that destroy phagosomal membranes and thus facilitate an efficient gene transfer from bacteria to the target cells^3–5^. The shRNA silencing TNF alpha was thus delivered to the target cells, which in turn significantly reduced the symptoms of inflammation in a widely used preclinical model of IBD.

## Results

In order to test whether the pSuper plasmid and their derivative psiTnfa vector, specifically designed to silence TNF alpha, actually penetrated into the distal colon cells the plasmid was engineered to incorporate a mouse pgk promoter driven *gfp* reporter whose expression was detected using specific PCR primers (Supplementary Figure 1.).

Dextran sulfate sodium (DSS)-induced colitis is one of the most commonly used models of ulcerative colitis^6^. It induces inflammatory infiltration, a reactive shortening of the colon and an increase in the expression of inflammatory mediators. In our experiments, we used two different concentrations, 2.5% and 3.5% DSS, to induce colon inflammation of two different levels of severity. As expected, both concentrations of DSS induced a significant increase in the inflammation score especially in the distal colon (Figure 1 A, B and Figure 2A, B). Similarly, the ulceration score also increased upon these treatments, with a more prominent effect in the distal colon (Figure 1 C, D and Figure 2 C, D). DSS induced inflammation is usually accompanied by a reactive shortening of the colon. Both concentrations induced a significant, approximately 30-35% shortening of the colon, (Figure 3 and Figure 4). Finally, in order to investigate the role of inflammatory mediators we measured their expression level in the experimental IBD model. 3.5% DSS induced a 2-fold induction of TNF alpha, a 100-fold induction of IL1a, an approximately 150 fold increase of IL1b and a 1.4 fold increase of IL6 (Figure 5).

**Figure 1:**
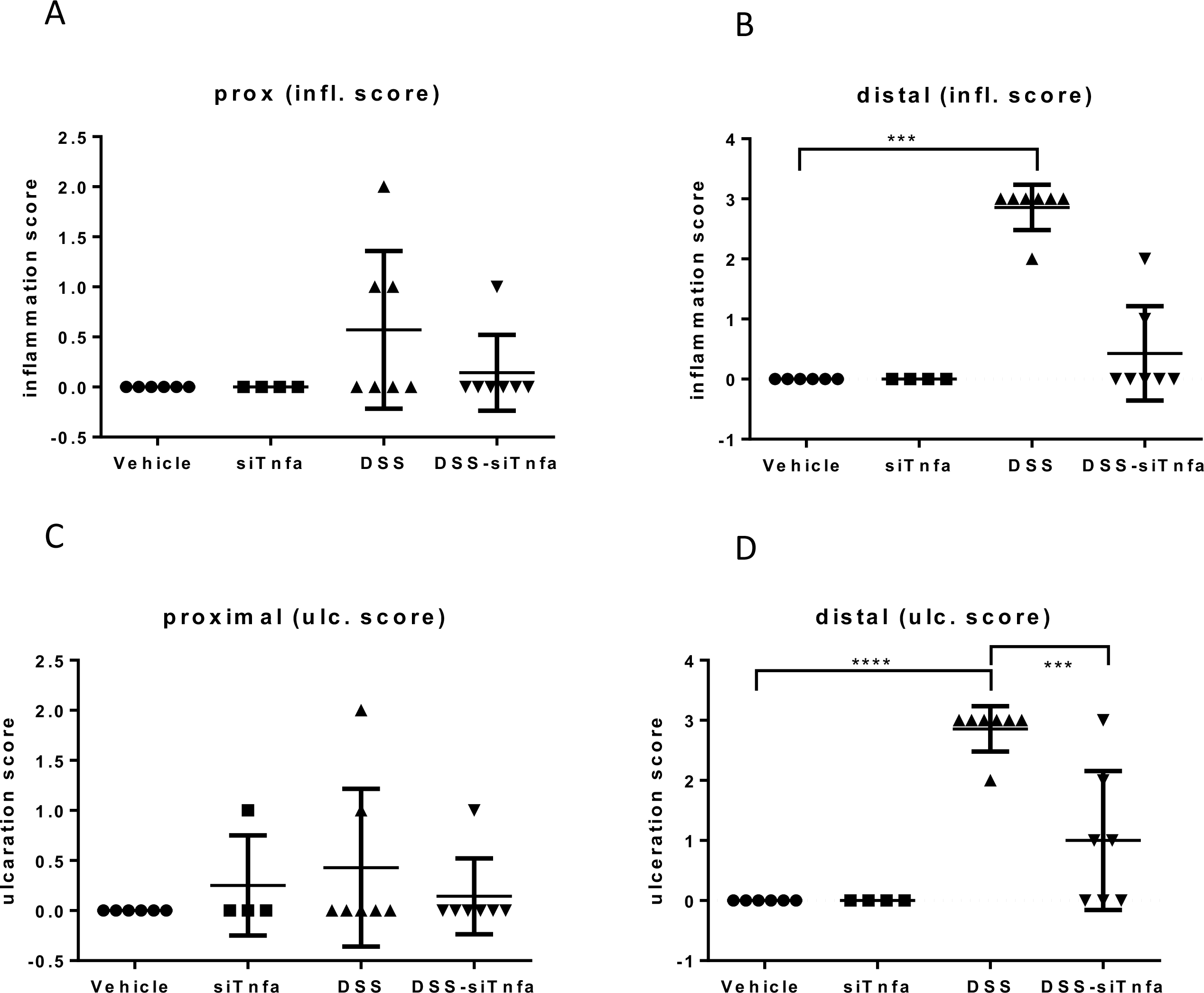
The colon inflammation and ulceration scores of the 2,5% DSS administered animals. Histological analysis was carried out on FFPE colon samples from experimental groups and the levels of inflammation and ulceration was scored as described in the methods. A: inflammation score of the proximal colon samples, B: inflammation score of the distal colon samples, C: ulceration scores of the proximal colon samples, D: ulceration scores of the distal colon samples. Vehicle: Tap water administered animal group, siTnfa: animals treated by modified *E. coli* MDS42 strains containing psiTnfa and invasive plasmids, DSS: colon specimen of the 2,5% DSS treated animals, DSS-siTnfa: animals administered 2,5% DSS and treated by modified *E. coli* MDS42 strains containing psiTnfa and invasive plasmids. Data represent the mean (SD). \*\*\**P* < 0.001, **** *P* < 0.0001.

**Figure 2:**
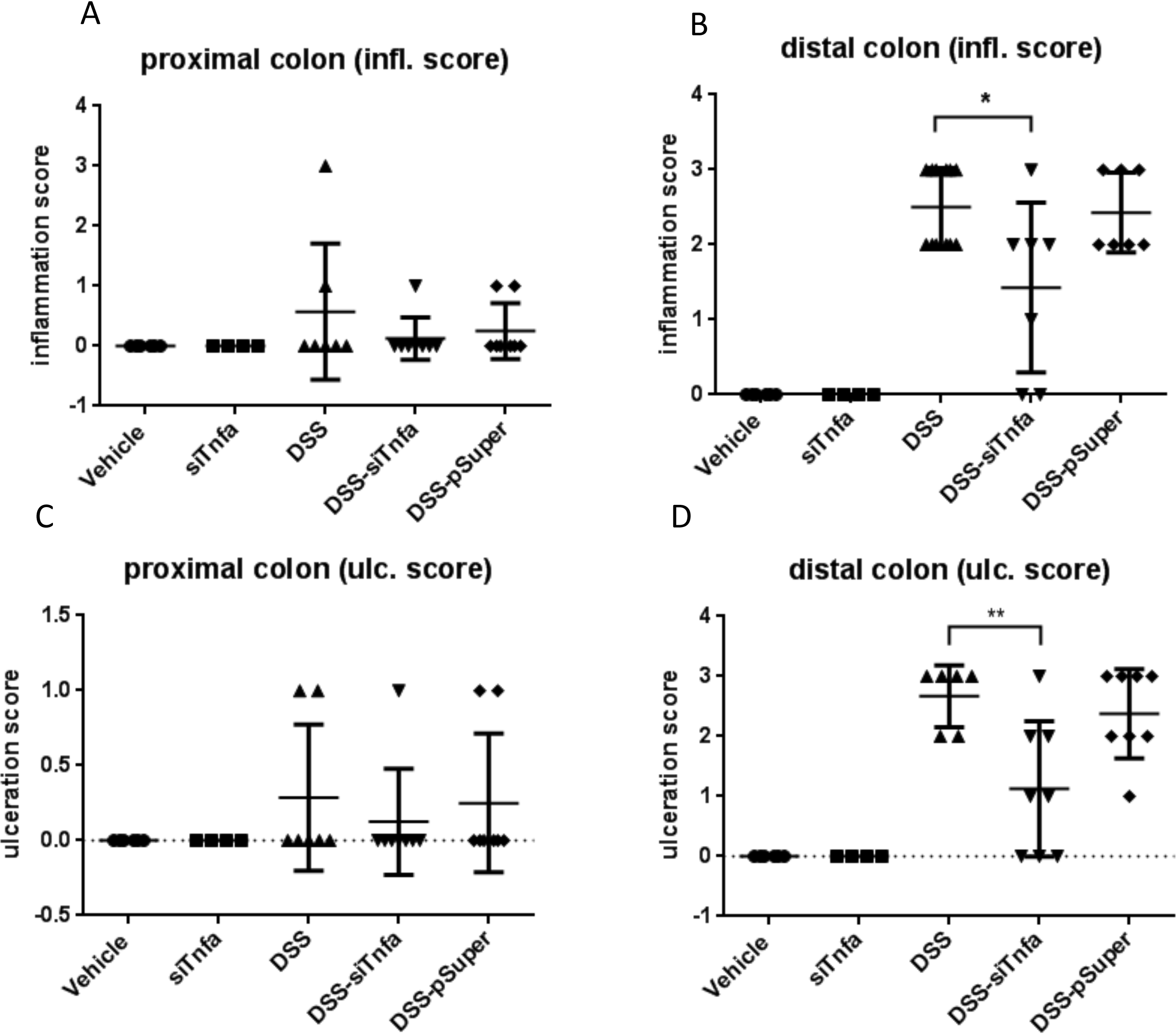
The colon inflammation and ulceration scores of the 3,5% DSS administered animals. Histological analysis was carried out on FFPE colon samples from experimental groups and the levels of inflammation and ulceration was scored as described in the methods. A: inflammation score of the proximal colon samples, B: inflammation score of the distal colon samples, C: ulceration scores of the proximal colon samples, D: ulceration scores of the distal colon samples. Vehicle: Tap water administered animal group, siTnfa: animals treated by modified *E. coli* MDS42 strains containing psiTnfa and invasive plasmids, DSS: colon specimen of the 3,5% DSS treated animals, DSS-siTnfa: animals administered 3,5% DSS and treated modified *E. coli* MDS42 strains containing psiTnfa and invasive plasmids, DSS-pSuper: animals administered 2,5% DSS and treated by modified *E. coli* MDS42 strains containing pSuper (empty vector) and invasive plasmids. Data represent the mean (SD). \**P* < 0.05, ** *P* < 0.01.

**Figure 3:**
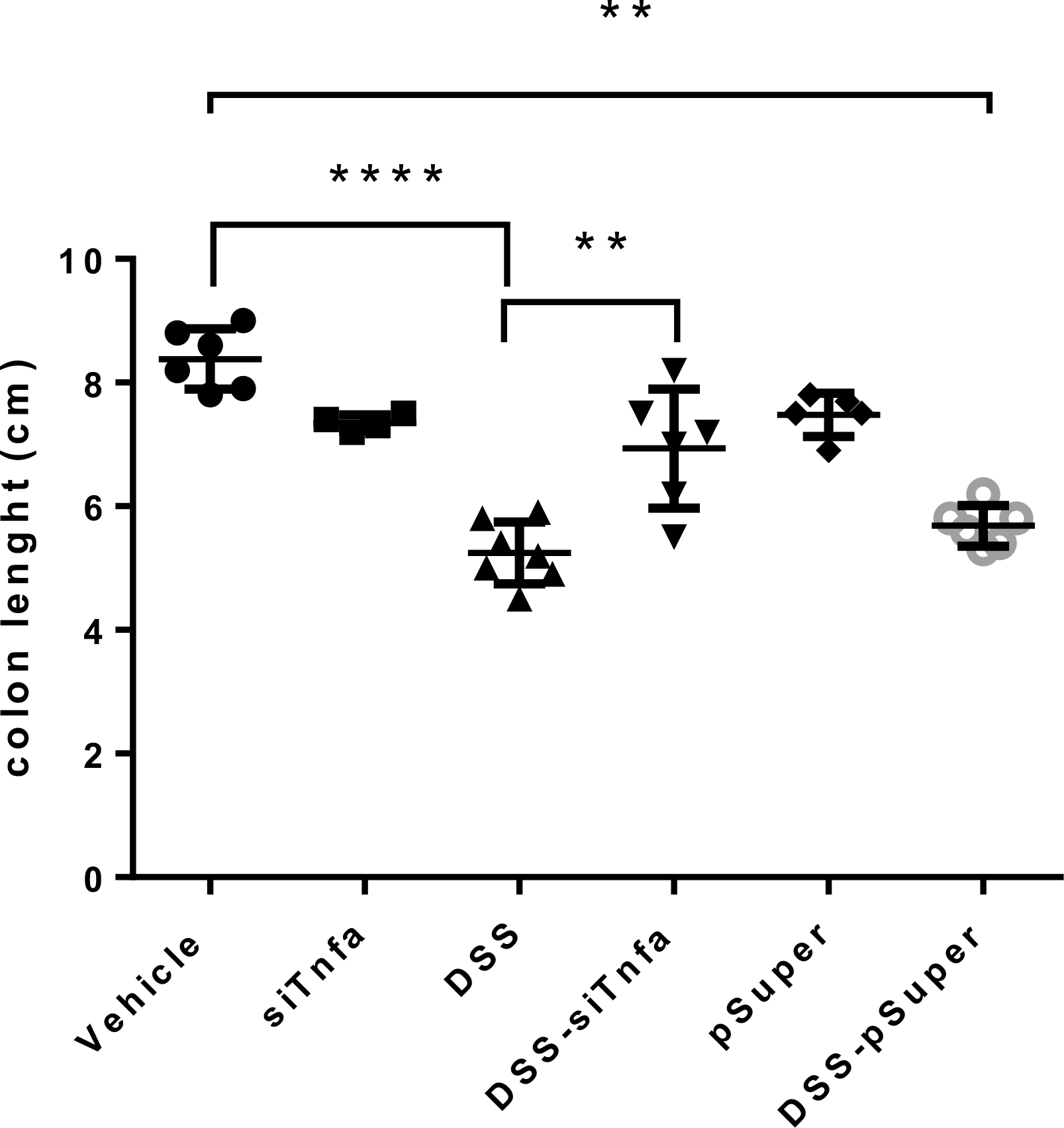
Effect of the modified *E. coli* MDS42 strains containing psiTnfa and invasive plasmids on the colon length. Vehicle: Tap water administered animal group, siTnfa: animals treated by modified *E. coli* MDS42 strains containing psiTnfa and invasive plasmids, DSS: colon specimen of the 2,5% DSS treated animals, DSS-siTnfa: animals administered 2,5% DSS and treated by modified *E. coli* MDS42 strains containing psiTnfa and invasive plasmids, pSuper: animals treated by modified *E. coli* MDS42 strains containing pSuper (empty vector) and invasive plasmids, DSS-pSuper: animals administered 2,5% DSS and treated by modified *E. coli* MDS42 strains containing pSuper (empty vector) and invasive plasmids. Data represent the mean (SD). \*\**P* < 0.01, **** *P* < 0.0001.

**Figure 4:**
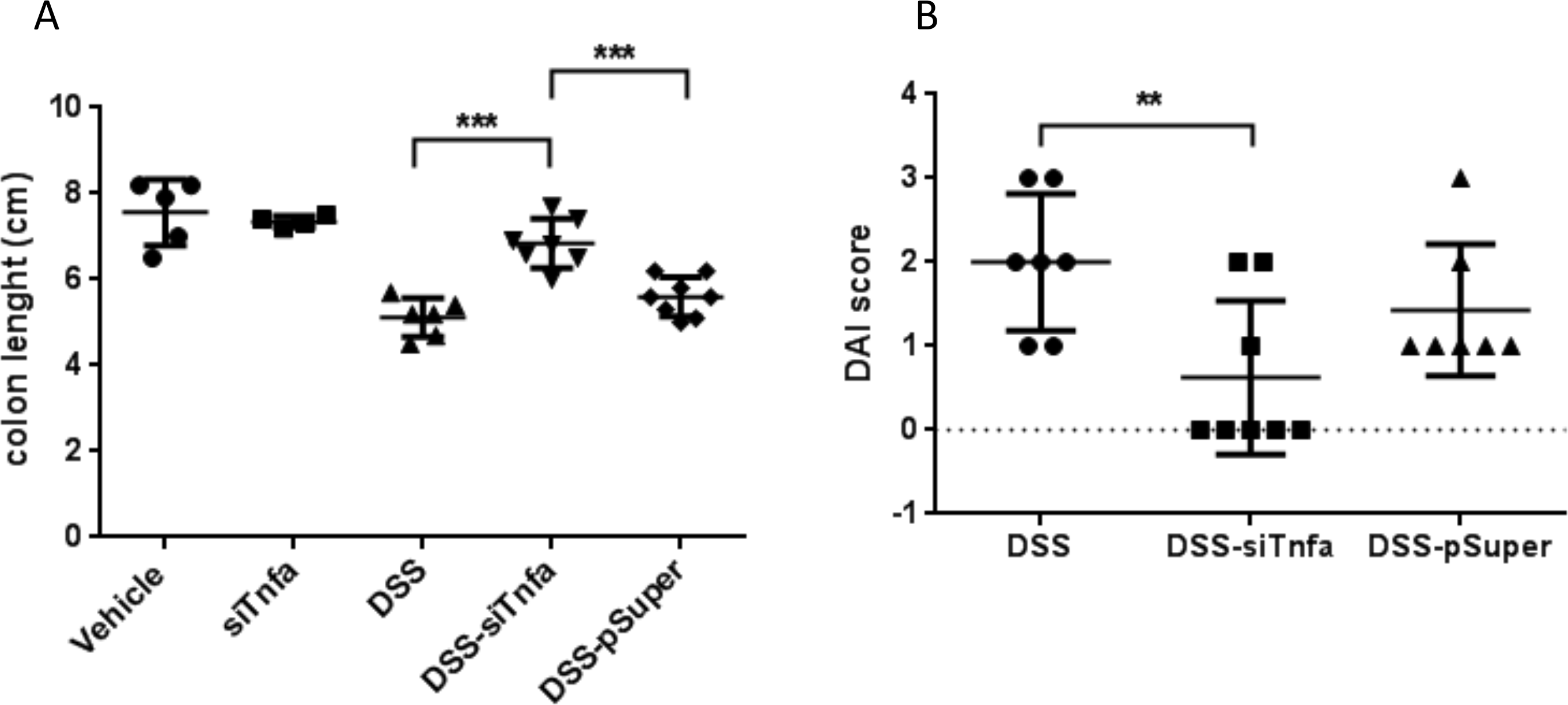
The effect of the 3,5% DSS induced colitis on colon length and the disease activity index (DAI) of colitis. A: Effect of the modified *E. coli* MDS42 strains containing psiTnfa and invasive plasmids on the colon length. The animals administered 3,5% DSS. B: The biological effect of modified *E. coli* MDS42 strains containing psiTnfa and invasive plasmids were evaluated considering the disease activity index (DAI) of colitis. Vehicle: Tap water administered animal group, siTnfa: animals treated by modified *E. coli* MDS42 strains containing psiTnfa and invasive plasmids, DSS: colon specimen of the 2,5% DSS treated animals, DSS-siTnfa: animals administered 2,5% DSS and treated by modified *E. coli* MDS42 strains containing psiTnfa and invasive plasmids, DSS-pSuper: animals administered 2,5% DSS and treated by modified *E. coli* MDS42 strains containing pSuper (empty vector) and invasive plasmids. Data represent the mean (SD). \*\**P* < 0.01, *** *P* < 0.001.

**Figure 5:**
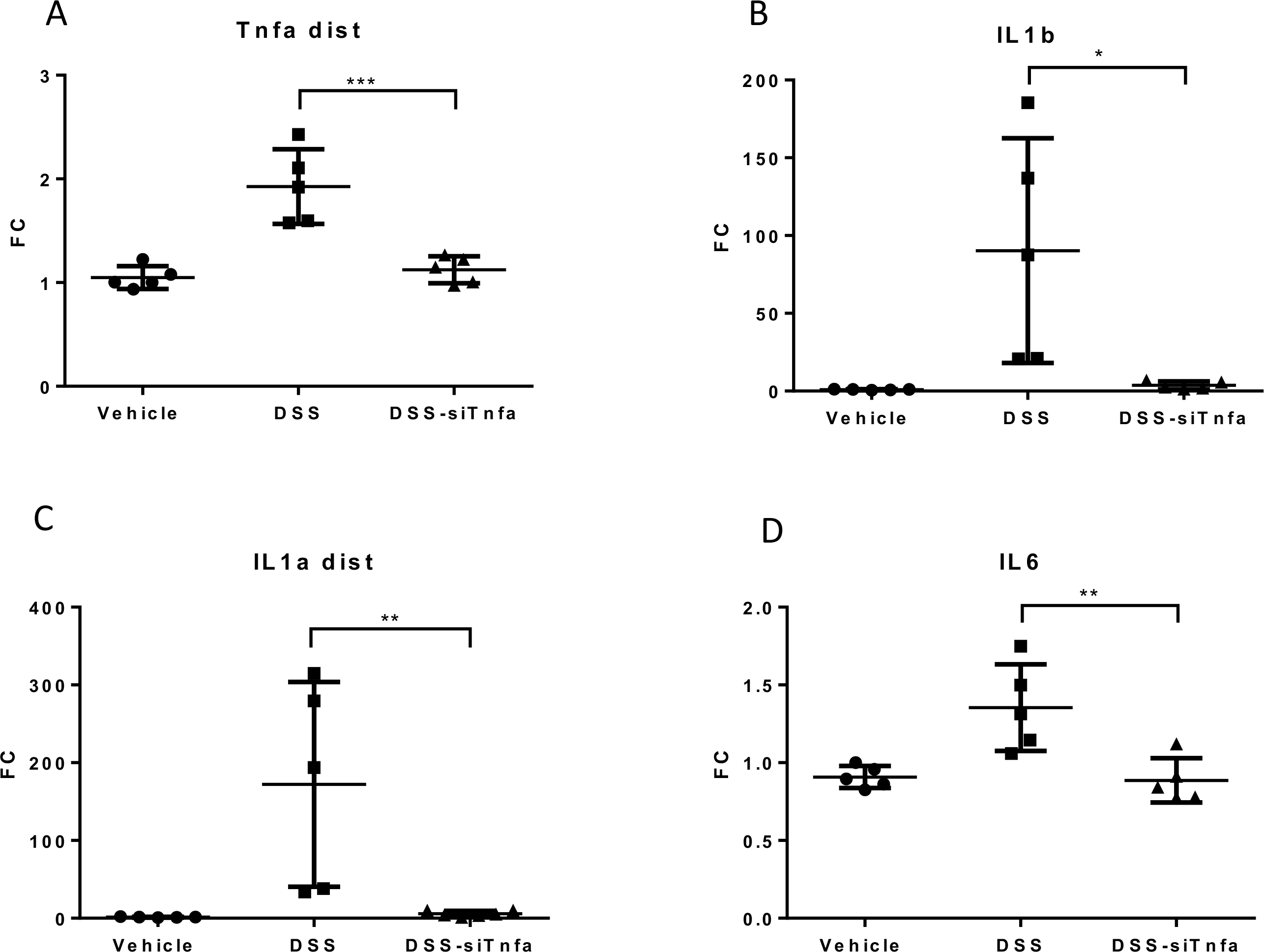
mRNA expression of proinflammatory cytokines on 3,5% DSS induced colitis. Colitis and the modified *E. coli* MDS42 strains containing psiTnfa and invasive plasmid treatment induced proinflammatory cytokines mRNA levels were measured by qPCR. A: Tnfa mRNA expression measured on distal colon samples, B: Il-1b mRNA expression measured on distal colon samples, C: Il-1a mRNA expression measured on distal colon samples, D: Il-6 mRNA expression measured on distal colon samples. Vehicle: Tap water administered animal group, DSS: colon specimen of the 3,5% DSS treated animals, DSS-siTnfa: animals administered 3,5% DSS and treated by modified *E. coli* MDS42 strains containing psiTnfa and invasive plasmids. Data represent the mean (SD). \**P* < 0.05, ** *P* < 0.01, *** *P* < 0.001.

The animals exposed to the induced IBD were then administered the *E.coli* MDS42 bacteria specifically engineered to silence TNF alpha. This treatment had a dramatic effect on the DSS-induced colitis. The increase of inflammatory mediators was significantly reduced. For each of the inflammatory mediators measured the expression returned to baseline levels. (Figure 5). Consequently, the inflammation score was also significantly reduced. In the case of 2.5% DSS treatment the inflammation score returned to baseline or near baseline levels both in the proximal and distal colon. The ulceration score increase was also reduced by 64% (Figure 1). For the higher DSS concentration (3.5%), both the inflammation and ulceration score were reduced by 50% in the distal colon (Figure 2). Finally, the colon length shortening was also significantly reduced to around the level seen in the control animals (Figure 3).

As a control we engineered the same bacteria carrying an empty vector, without the shRNA for TNF alpha. This treatment did not reduce the various measures of inflammation. Similarly, the colon length of the animals treated with only pSuper and psiTnfa plasmids (with invasion plasmid) showed no significant alteration in comparison with the control (water only) (Figure 3 and Figure 4). The mucosa thickness did not show any significant change in the proximal or distal colon regions for any of the above listed treatments (data not shown).

For 3.5% DSS treatment we also determined the DAI (disease activity index) value indicating the severity of the inflammatory processes. Again, this was significantly reduced by the engineered bacteria delivering the Tnfa-specific shRNA responsible for Tnfa silencing (Figure 4B).

## Discussion

During the past decade, engineered bacteria have emerged as a potential “living” vehicle delivering both diagnostic and therapeutic molecules to the gastrointestinal tract^7^. There are several approaches to the therapeutic exploitation of such a biological system in terms of how directly the bacteria are interacting with the host biological system. For example, bacteria were engineered to express phenylalanine metabolizing enzymes that, upon administration to the digestive tract, would reduce the symptoms of phenylketonuria^8^. In this case, the bacteria exert their therapeutic effect inside the colon lumen by breaking down the phenylalanine introduced as part of the diet. Similarly, lactic acid bacteria exerted their therapeutic effect by producing elafin inside the gut, and thus reducing the inflammation in the same preclinical model that we used in this work^9^. In these cases, both the bacteria and the therapeutic effectors produced by them remained in the intestinal lumen. Considering the often aggressive and therapy resistant nature of inflammatory bowel disease^10^ we decided to test a more direct approach and deliver shRNA silencing of one of the key pathogenic inflammatory mediators, TNF alpha, directly into the target cell. A similar strategy was previously tested by *E. coli* expressing invasin, listeriolysin O cytolisyn and specific shRNA that could silence the β-1 catenin gene in the colon of experimental animals^11^. We adjusted this approach to target one of the key mediators of IBD, TNF alpha. Our results suggest, that this direct delivery of therapeutic modulators can cause a significant reduction of inflammation.

There are serious safety concerns about the use of live bacteria in the therapeutic setting. Bacteria may exchange genetic material with other bacteria in the colon, which may lead to uncontrollable or difficult to control consequences^12^. This risk could potentially be prevented by strategies that would introduce mechanisms into the bacterial vectors leading to their self-destruction, such as the toxin-antitoxin systems^13^, within a few days in the colon but still able to deliver the therapeutic effect.

## Materials and Methods

### Animals

Adult (8-10 weeks old) male FVB/Ant mice were obtained from the local colony bred at the Medical Gene Technology Unit (Specific Pathogen Free, SPF level) at the Institute of Experimental Medicine Budapest Hungary. Animals were housed at the Minimal Disease (MD) level, 3-5/cage under controlled environmental conditions: temperature, 21°C±1° C; humidity, 65%; light-dark cycle, 12-h light/12-h dark cycle, lights on at 07:00. Mice had free access to rodent food and water. All procedures were conducted in accordance with the guidelines set by the European Communities Council (86/609/EEC/2 and 2010/63 Directives of European Community) and the protocol was approved by the Institutional Animal Care and Use Committee of the Institute of Experimental Medicine, Budapest Hungary (permit number: PEI/001/29-4/2013).

### UC model generation and animal treatment

In animal models, UC can be induced by DSS administration. The experimental animals were 10 weeks old Fvb / Ant male mice. Mice were treated with drinking water mixed 2.5% and 3.5% DSS, which was administered for 8 days. The engineered bacterial cells were grown in LB liquid medium at 37 °C. The overnight bacterial culture was harvested by centrifugation and washed twice with sterile tap water to eliminate the antibiotic contamination (ampicillin and spectinomicin). Finally, the cell pellet was suspended in sterile tap water and administered to the animals. The engineered bacterial cells (2-5x 10^5^ CFU) and vehicle (sterile tap water) was administered daily by oral gavages. During treatment, DAI (Disease Activity Index) values were observed. At the end of the experiment, the animals were anesthetized and decapitated. Colon samples were frozen and fixed for histological processing. Intestinal contents were frozen.

### Cloning procedures

The nucleotide sequences encoding shRNS for efficient silencing the Tnfa was designed using the Serial Cloner 2.6.1 software. The forward and reverse synthetic DNA oligos (Microsynth, Germany) were heated in annealing buffer (100 mM NaCl, 50 mM Hepes, pH 7.4) at 90 ° C for 5 minutes and allowed to cool to room temperature. The annealed double stranded synthetic DNA oligos were ligated into pSuper retro neo GFP (Oligoengine, USA) vector after linearisation with BglII and HindIII enzymes. During the cloning steps, the ligation was heated to 90 ° C for 5 minutes and allowed to cool to room temperature. Subsequently, conventional heat shock transformation was introduced into the prepared *E. coli* MDS42 strain. Recombinant bacteria were selected on LB medium containing ampicillin. Plasmid constructs encoding the resulting psiTNFα (shRNA) precursors were verified by nucleotide sequence assay and restriction enzyme digestion. Using the High-Speed Plasmid Mini kit (Genaid), a plasmid was isolated from recombinant colonies and digested with EcoRI and HindIII enzymes. The digested plasmids were checked by agarose gel electrophoresis (1% agarose (Sigma), 1xTris-boric acid EDTA buffer, pH 8.5). The empty vector carries 1000 bp long insert, while the plasmids containing the ligated synthetic oligos are 281 bp long inserts resulted after double digestion. Plasmids from positive colonies were analyzed by Sanger’s nucleotide assay using a specific sequencing primer (5’-ggaagccttggcttttg-3 ’). The *invazin* gene of the pGB2-Ω plasmid was verified by PCR amplification and restriction enzymatic digestion PstI. PstI removes the fragment from the plasmid that contains the *invasin* and *histeriolysin*.

### Quantitative Real-Time PCR

Frozen tissue samples were homogenized in TRI Reagent Solution (Ambion, USA) and total RNA was isolated with QIAGEN RNeasy Mini Kit (Qiagen, Valencia, CA, USA) according the manufacturer’s instruction. To eliminate genomic DNA contamination DNase I treatment were used and 100 μl Rnase-free DNase I (1 unit DNase) (Thermo Scientific, USA) solution was added. Sample quality control and the quantitative analysis were carried out by NanoDrop 2000 (Thermo Scientific, USA). Amplification was not detected in the RT-minus controls. The cDNA synthesis was performed with the High Capacity cDNA Reverse Transcription Kit (Applied Biosystems, USA). Primers for the comparative Ct experiments were designed by Primer Express 3.0 Program and Primer Blast software. The primers (Microsynth, Balgach) were used in the Real-Time PCR reaction with Fast EvaGreen® qPCR Master Mix (Biotium, USA) on ABI StepOnePlus (Applied Biosystems, USA) instrument. The primer sequences are listed Supplementary Table 1. The gene expression was analyzed by ABI Step One 2.3 program. The amplicon was tested by Melt Curve Analysis on ABI StepOne Plus instrument. Experiments were normalized to *gapdh* expressions.

### GFP tracing in the colon

Total RNA was isolated and cDNA synthesis was performed from the frozen colon samples. The *gfp* expression in the colon tissue was demonstrated by PCR using specific oligonucleotid primers (for: GGA CGA CGG CAA CTA CAA GA, rev: AAG TCG ATG CCC TTC AGC TC). The PCR product was separated and stained on agarose gel.

### Histology

Formalin fixed colon samples were processed and embedded in paraffin using the standard protocol. Sections of 4 μm were stained with haematoxylin and eosin. Slides were analyzed by an expert pathologist in a blinded manner.

### Statistical analysis

Data are expressed as means ± SD. The data were first subjected to a Kolmogorov-Smirnov normality test. Data passing this test were analyzed by One-way ANOVA followed by the Tukey’s *post hoc* test. If the data showed non-Gaussian distribution, the Kruskal-Wallis test was used. Statistical analysis was performed using GraphPad PRISM version 6 software (GraphPad Software, USA). *P* ≤ 0.05 was considered significant.

## Acknowledgements

This work was supported by the National Research Development and Innovation Office Hungary (grant number: KFI_16-1-2017-0245).

---

**Supplementary Figure 1:**
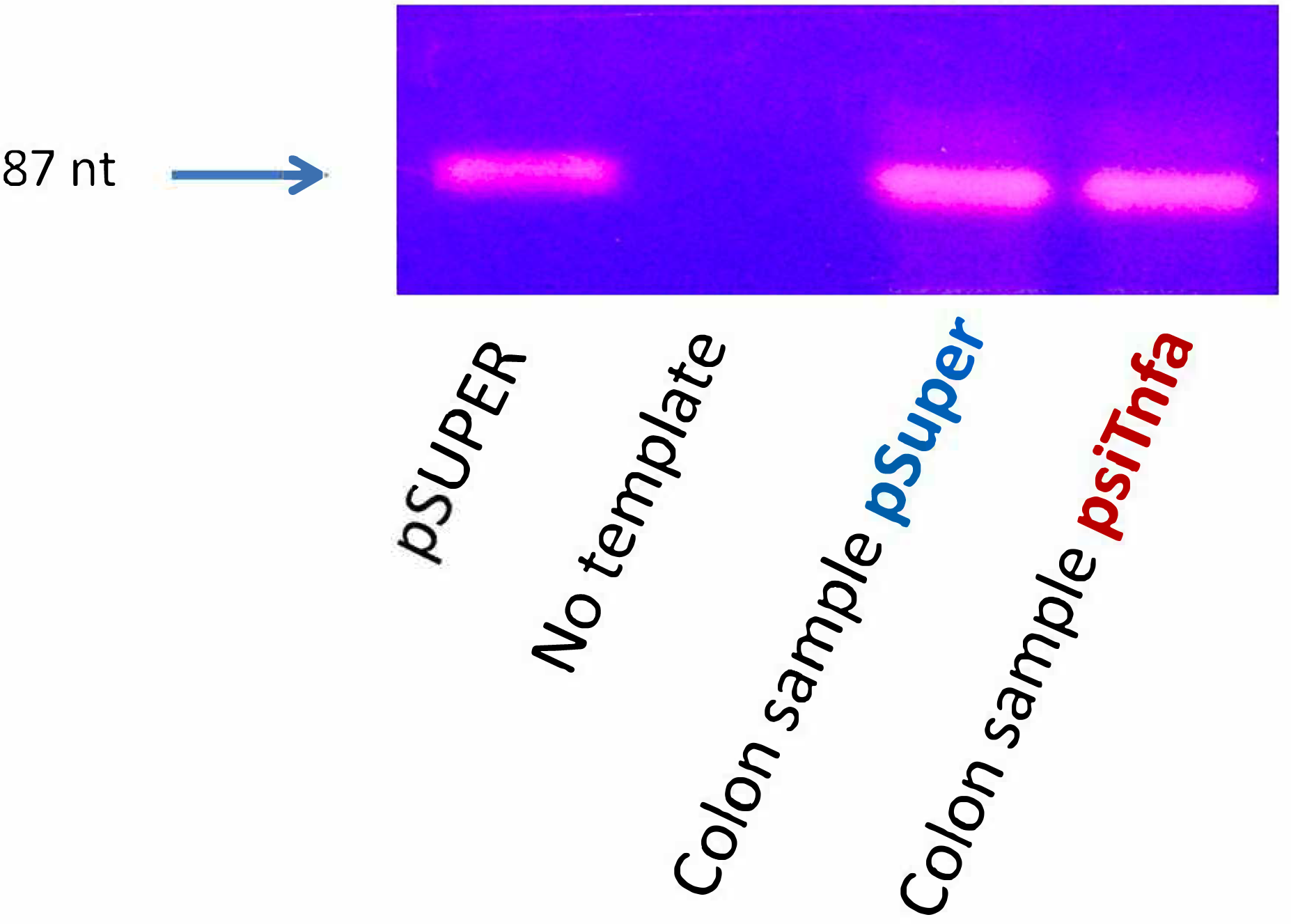
Total RNA was isolated and cDNA synthesis was performed from the distal colon samples. The *gfp* expression in the colon tissue was demonstrated by PCR using *gfp* specific oligonucleotid primers. The PCR product was separated and stained on 2% agarose gel.

**SM Table 1.:**
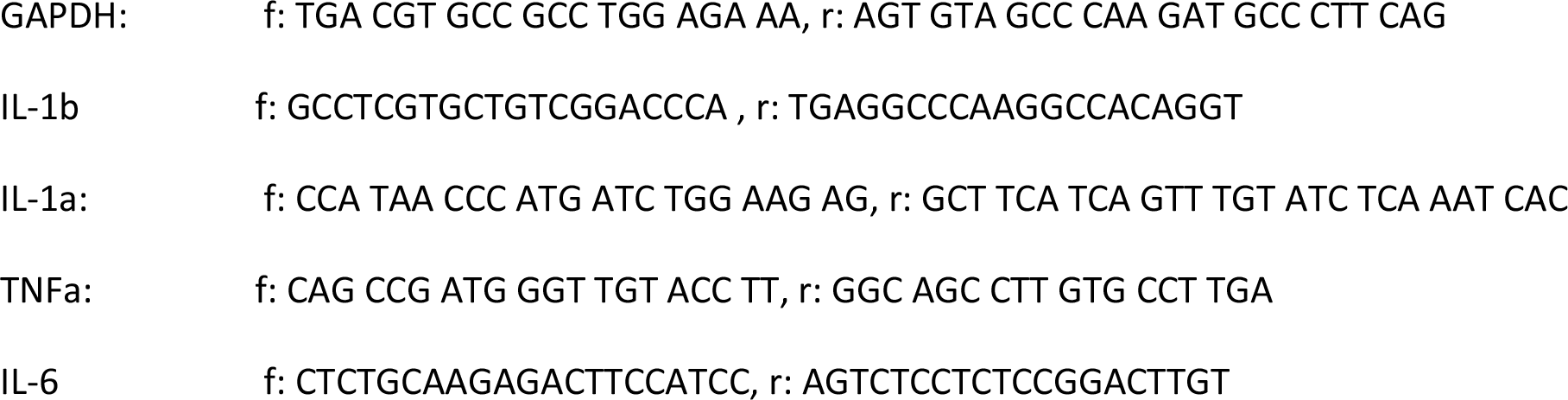
Primer sequences used in qRT PCR reactions.

